# Novel Functional metabolites that affect biofilm formation are regulated by bioavailable iron with siderophore-dependent pathway

**DOI:** 10.1101/2020.03.04.977660

**Authors:** Rui Guo, Xilin Luo, Jingjing Liu, Haitao Lu

**Affiliations:** Key Laboratory of Systems Biomedicine (Ministry of Education), Shanghai Center for Systems Biomedicine, Shanghai Jiao Tong University, Shanghai 200240, China; Laboratory for Functional Metabolomics Science, Shanghai Jiao Tong University, Shanghai 200240, China

**Keywords:** Biofilm Formation, Iron Bioavailability, Functional Metabolites, Siderophore Biosynthesis, Targeted Metabolomics

## Abstract

Biofilms are broadly formed by diverse microorganisms under stressful environments and are basically surrounded by an EPS matrix, enabling bacterial cells to confer more resistance to biocides, antibiotics and other invasions than their planktonic counterparts. However, biofilm formation causes problems in various fields, including clinical infections, environmental pollution, agricultural production and industrial contamination. Unfortunately, the mechanism of biofilm formation has not been completely elucidated, and currently, we lack an efficient strategy to tackle these tough problems and destroy biofilms. In the present study, we sought to decipher the mechanism of biofilm formation through the regulation of functional metabolites regulated by iron. By exposing bacterial cells to various concentrations of iron, we found that iron can regulate biofilm formation, and phenotypic changes were obviously dependent on iron concentration. A functional metabolome assay was further implemented to investigate the regulatory mechanism of iron on biofilm formation; we verified that siderophores (linear enterobactin, yersiniabactin, di-glucosylated-salmochelin and HPTT-COOH) mostly account for the transportation of iron into bacterial cells. Then, bioavailable iron was recruited by bacterial cells to direct the biosynthesis and expression of five functional metabolites (L-tryptophan, 5’-MTA, spermidine, CMP and L-leucine), which were identified as new effectors that directly regulate biofilm formation. Taken together, this study is the first to identify five new metabolic effectors to efficiently regulate biofilm formation, the biosynthesis and expression of these functional metabolites can be targeted to tackle the challenging problems associated with biofilm formation in different fields.

## Introduction

Biofilms are self-protective, structurally organized communities that are broadly formed by diverse microorganisms under different stressful environments and can adhere to surfaces or liquid-air interfaces. The compositions of biofilms have been analyzed, and biofilms are basically surrounded by an extracellular polymeric substance (EPS) matrix consisting of nucleotides, proteins and extracellular polysaccharides; these components enable biofilms to confer more resistance against biocides, antibiotics and other invasive compounds than their planktonic counterparts^1–3^. Biofilms unexpectedly trigger outbreaks of challenging problems in different fields; for example, biofilms can lower agricultural production, decrease food safety, cause industrial contamination, and even lead to antibiotic resistance in biomedical sciences ^3–5^. Although the scientific community has made great efforts in understanding the biochemistry of biofilm formation over past decades, the mechanism of biofilm formation remains incompletely understood. However, the available strategies to solve these problems and destroy biofilms in different fields are insufficient. Surprisingly, our recent study illustrated that substantial metabolic reprogramming occurs during biofilm formation^6^; thus, we maintain that metabolism is a new window allowing us to better understand the mechanism of biofilm formation and enabling us to explore the dissociation mechanism to prevent and/or reverse biofilm formation in the real world. In the present study, the uropathogenic strain *Escherichia coli* (*E. coli*) UTI89 was used as an experimental organism, and *E. coli* UTI89 was shown to often form stable biofilms during infection^7–10^, leading to the need for high-frequency antibiotics and recurrence in the clinic. Furthermore, we combined structural imaging with targeted metabolomics methods to comprehensively capture the functional features of biofilm formation, visualize the structure-phenotype and decipher the mechanism of metabolic regulation. Compared to previous studies, our data verify the mechanism of metabolic regulation during biofilm formation, which were phenotypically validated with structural imaging observations. Generally, the selected siderophores can transport iron into bacterial cells, and iron is recruited to regulate the biosynthesis and expression of five functional metabolites, including L-tryptophan, 5’-MTA, spermidine, CMP and L-leucine, which then regulate biofilm formation in different patterns. Taken together, this study is the first to elucidate the mechanism of biofilm formation through siderophores that affect the bioavailable iron, which regulates functional metabolites; the results of this study will largely assist in developing biofilm clearance-based strategies and answering the difficult questions regarding biofilm formation in different niches.

## Materials and methods

### Chemicals and reagents

Distilled water was purchased from Watsons Corporation (Guangzhou, China). Acetonitrile, methanol, and formic acid (HPLC grade) were purchased from Fisher Scientific (Shanghai, China). LB broth (Luria−Bertani) and LB agar were purchased from Becton Dickinson (Sparks, USA). Casamino acid, yeast extract, magnesium sulfate, manganese chloride, L-aspartate, putrescine, L-isoleucine, L-histidine, CMP, L-tryptophan, L-leucine, spermidine and 5’-MTA were of USP grade and were purchased from Sigma-Aldrich Corporation (Saint Louis, USA). All other reagents were of ACS grade.

### Bacterial strains and cultivation

To trigger biofilm formation under experimental conditions, a static culture system was prepared with colony-forming antigen (CFA) medium (1% casamino acids, 0.15% yeast extract, 0.005% magnesium sulfate and 0.0005% manganese chloride). The protocol for biofilm formation was briefly summarized as follows: after 12 h of culture on LB-agar plates, one colony of the UTI89 strain was incubated in LB broth for 4 h then further diluted into CFA medium at a ratio of 1:1000 to prepare cultures with different concentrations. All the cultures were statically cultivated in flask incubators for 72 h at 30°C to trigger mature biofilm formation. For the cultures treated with FeCl_3_ and the functional experiment, all the analytical compounds were first diluted and added to the CFA medium at the defined concentration; then, the remaining culture procedure was the exact same as the protocol mentioned above.

### Sample preparation

After 72 hours of culture, biofilms were isolated from 50 ml culture solutions, and the suspensions in the culture medium were centrifuged at 2000 rpm and 4°C for 15 min to collect the planktonic cells. The biofilms were washed with 2 ml of PBS 3 times; then, 2 ml of 80% ice-cold methanol was added, and the solution was homogenized on ice for 2 min. This procedure was repeated three times. Next, the samples were centrifuged at 12000 rpm at 4°C for 15 min, and the supernatants were mixed with 800 μl of ice-cold acetonitrile for 20 min before centrifugation at 12000 rpm at 4°C for 15 min. Finally, the supernatants were passed through a 0.22 μm filter membrane (nylon) before lyophilization. The lyophilized samples were stored at −80°C for further experimental use.

### Siderophore extraction

The extraction protocol was modified from previous publications (12, 13). Briefly, 12 μL of 0.1 M FeCl_3_ was added to 2 mL of biofilm supernatant, and the mixture was filtered with a 0.22 μm filter membrane (nylon). After 15 min incubation at room temperature, centrifugation was performed to collect the supernatant, which was uploaded into an SPE column (Waters, USA) containing 60 mg of Oasis HLB sorbent. Each column was washed with 0.5 mL of 5% methanol once, and then siderophores were eluted with 0.5 mL of 100% methanol. Finally, 5 μL of elute was injected into a LC-QTOF/MS for the siderophore assay.

### Targeted metabolomics method

For the targeted metabolomics assay, referring to our protocol (11), lyophilized samples were resuspended in 200 μL of water; 5 μL of the sample was used for targeted metabolomics analysis, which was performed with an UHPLC system (Agilent 1290 Infinity, Agilent Technologies, USA) coupled to a triple quadrupole (QQQ) mass spectrometer (Agilent 6495 QQQ, Agilent Technologies, USA). RPLC separation was performed with an ACQUITY UPLC HSS T3 (2.1×100 mm, 1.8 μm) column; mobile phase A was water with 0.1% formic acid (v/v), and mobile phase B was acetonitrile with 0.1% formic acid (v/v) with an optimized gradient-elution program: 0-2 min, 98% A; 2-10 min, 98%-65% A; 10-12 min, 65%-20% A; 12-14 min, 20%-2% A; 14-30 min, 2% A. HILIC separation was carried out with an ACQUITY UPLC BEH amide column (2.1 mm i.d×100 mm, 1.7 μm; Waters); mobile phase A was changed to water with 0.1% (v/v) formic acid and 10 mM ammonium acetate, while mobile phase B was acetonitrile with 0.1% formic acid (v/v). The relevant gradient-elution program was optimized as follows: 0-4 min, 5%-12% A; 4-15 min, 12%-50%; 15-25 min, 50% A. The flow rate for both RPLC and HILIC separation systems was set at 0.3 mL/min, and the column temperature was maintained at 40°C. All the samples were placed at 10°C with a 5 μL-volume injection. The parameters of MS acquisition were summarized as follows: sheath gas temperature, 380°C; sheath gas flow, 12 L/min; dry gas temperature, 250°C; dry gas flow, 16 L/min; capillary voltage, 4000 V in positive mode and 3500 V in negative mode; nozzle voltage, 1500 V; and nebulizer pressure, 20 psi; the acquisition time was set at 14 min. The dynamic MRM parameters for targeted metabolomics involving 240 metabolites have been recorded in our preprint publication ^11^.

### Siderophore assay

For the siderophore assay, the samples were analyzed sequentially with an UHPLC (Agilent 1290 II Infinity, Agilent Technologies, USA) coupled to an IM-QTOF mass spectrometer (Agilent 6560 IM-QTOF, Agilent Technologies, USA) system. The column used for sample separation, injection volume, flow rate, column temperature and mobile phases A and B were the same as the targeted metabolomics method listed above. The gradient-elution program is summarized as follows: 0−1 min, 2% B; 16 min, 35% B; 20-37 min, 98% B; 38-50 min, 2% B. The analytical parameters of MS acquisition are listed as follows: sheath gas temperature, 350°C; sheath gas flow, 12 L/min; dry gas temperature, 300°C; dry gas flow, 10 L/min; capillary voltage, 4000 V in positive mode; nozzle voltage, 500 V; and nebulizer pressure, 40 psi. The acquisition time was set at 28 min, and the m/z for each siderophore was obtained from previous publications^12, 41^ (Extended Data Figure S4). Siderophores, including yersiniabactin, enterobactin, salmochelin, and HPTT-COOH, were determined with the developed method described above.

### CFU assay

Biofilms were dissected with a manual homogenizer in cold PBS to collect the free bacterial cells. The biofilm suspensions and planktonic cells were used to measure the CFU value. The following steps were performed according to a previous publication^13^.

### Scanning electron microscopy assay

The cells were washed with PBS 4 times; the specimens were dyed with 1% osmic acid solution and subsequently washed 4 times with PBS. Then, the mixtures were dehydrated by a gradient concentration of ethanol (50%, 70%, 90% and 100%) for approximately 10 min at each step. Finally, the dehydrated specimens were completely dried and analyzed by scanning electron microscopy (SEM) (TESCAN-MAIA3).

### Transmission electron microscopy assay

Mature biofilms and planktonic cells were fixed with 2.5% glutaraldehyde for 6 hours. The cells were washed with PBS four times; the specimens were dyed with 1% osmic acid solution and washed 4 times with PBS. Next, the samples were dehydrated by a gradient concentration of ethanol (50%, 70% and 90%), 90% ethanol:90% propanone=1:1, and 90% and 100% propanone for approximately 10 min each. After that, the cells were fixed with resin with propanone:epoxy ratios of 1:1 and 1:2 and fixed in a drying oven at 60°C for 48 h. Finally, all the specimens were sectioned with an ultramicrotome (LEICA EM UC7) and then analyzed by transmission electron microscopy (TEM) (Talos L120C G2).

### Crystal violet staining assay

Biofilm formation was quantified by crystal violet staining assay. The protocol was summarized as follows: the mature biofilm was washed 3 times with PBS, and then the biofilm was fixed with methanol for 15 min. After the methanol was volatilized, 100 μl of 0.01% crystal violet was added to the biofilm and incubated for 20 min. The dye wells with crystal violet were washed 5 times with PBS. Bound crystal violet was solubilized with 100 μl of 30% acetic acid for 30 min and then shaken slowly. The OD was measured with a microplate reader at 570 nm.

### Data analysis and visualization

The MS raw data generated from the biological samples were first processed by qualitative analysis with the in-house software Agilent Corps, which integrated the signal peaks and generated three-dimensional data matrixes with the peak area of the metabolite-metabolite ID-sample ID. After the peak area for each metabolite was normalized to the tissue weight of each sample, the processed data were uploaded onto MetaboAnalyst, a free online software (http://www.metaboanalyst.ca/MetaboAnalyst/), for partial least-squares discriminant analysis (PLS-DA) and heatmap vs hierarchical-cluster analysis (HCA). Bar plots and all other statistics were generated using GraphPad Prism 6.0, Microsoft Office (Excel 2013) and MetaboAnalyst version 4.0.

## Results

### Phenotype-image validates the organized structures of biofilms

To analyze the biofilm formation of the model organism *E. coli* UTI89, we employed imaging methods, such as TEM and SEM, to comprehensively capture the structural features of biofilms compared to their planktonic counterparts. The TEM images of two bacterial patterns were observed, and the biofilms had a denser layer of fibers involving the EPS component compared to the relevant planktonic populations (Figures 1a and d). In addition, SEM revealed that the bacterial cells embedded in the biofilm were loosely packed and surrounded by an extensive fibrous network, which was extremely different from the planktonic cells (Figures 1b, c, e and f). This observation is consistent with previous studies^6, 10^, and we could argue that biofilms are not randomly formed but have uniquely organized ultrastructures compared to planktonic cells. In addition, crystal violet and CFU (colony-forming units) assays were applied to quantitatively assess biofilm formation, and the data demonstrated substantial growth differences between biofilms and planktonic cells (Figure S1). In short, biofilms have a unique organized structure, and this was confirmed by the UTI89 strain, which was grown in defined culture medium. Such phenomena might be implicated in substantial metabolic reprogramming during biofilm formation^6^.

**Figure 1.**
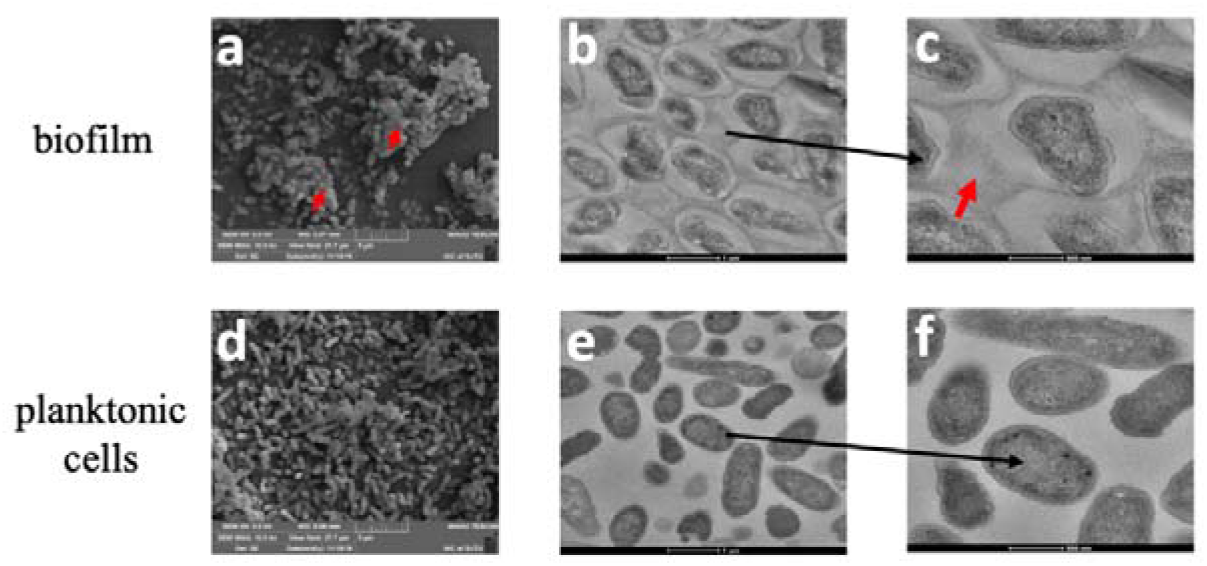
Phenotypic comparison revealed that biofilms have a unique organized structure compared to their planktonic counterparts. (a) SEM image of biofilm; (b and c) TEM image of biofilm. (d) SEM image of planktonic cells. (e and f) TEM image of planktonic cells.

### Biofilm formation has unique metabolic characteristics and are considerably differentiated from their planktonic counterparts

To further explore the metabolic characteristics of biofilm formation, a newly developed targeted metabolomics method^11^ was used to characterize the different cell metabolomes between biofilms and planktonic cells. To capture the precise metabolomes, we only collected metabolite data with stable and reproducible peaks to inspect the metabolic features of biofilm formation (Table S1). The precision-targeted metabolomics assay deciphers distinct metabolic characteristics of biofilm formation that are completely differentiated from those of the planktonic population (Figure 2). A total of 40 small-molecule metabolites were recognized that account for the unique metabolic characteristics of biofilms, and they were mostly part of many key metabolic pathways, such as carbohydrate metabolism, amino acid metabolism, and nucleotide metabolism (Table S2 **and** Figure S2). Some of the metabolites were reported in our previous study, including L-aspartate, dGMP, spermidine and L-isoleucine^6^. In addition, many new metabolites associated with biofilm formation were discovered and identified in this study. Our data reveal that pyridoxal 5’-phosphate was significantly increased during biofilm formation. Many L-amino acids or derivatives involving L-glutathione oxidized, N-acetyl-glutamic acid, L-tryptophan, L-glutamine, L-valine, L-arginine, L-proline, L-histidine, L-ornithine, and L-leucine were markedly altered during biofilm formation, supporting our previous finding that L-amino acids mainly participate in biofilm formation^6^.

**Figure 2.**
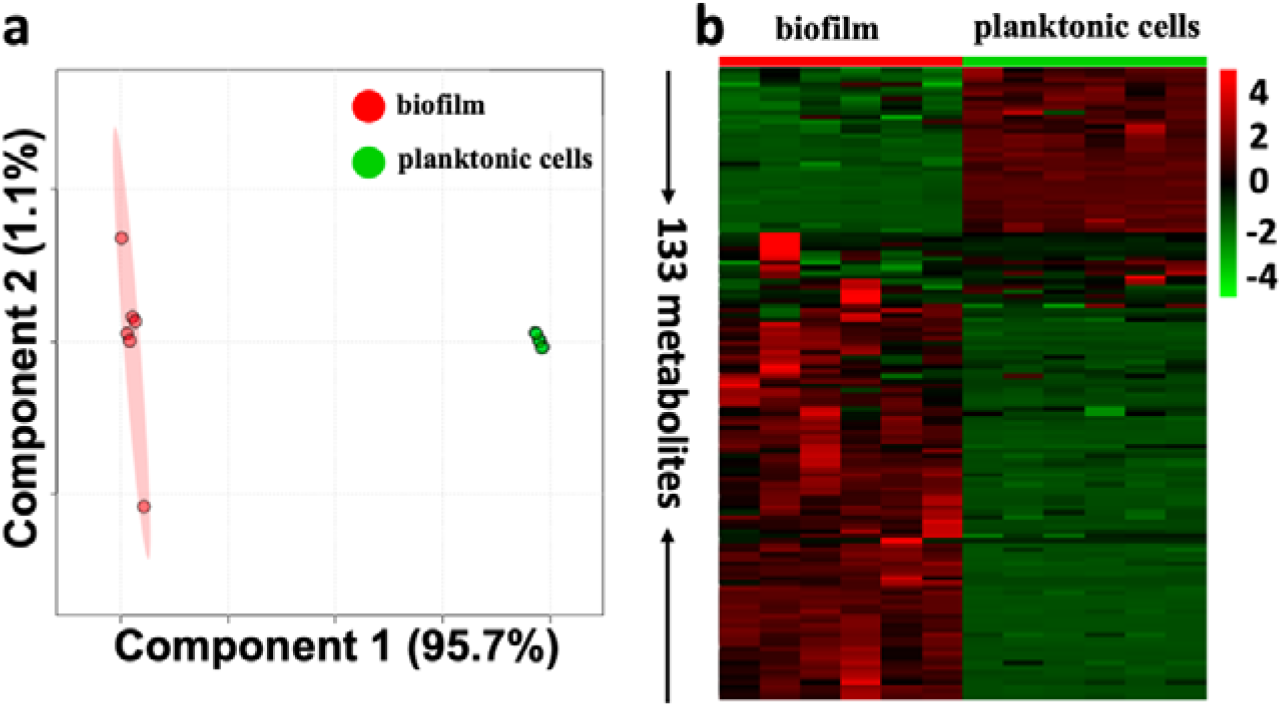
A precision-targeted metabolome assay revealed distinct metabolism that can obviously distinguish biofilms from planktonic cells. (a) Score plot resulting from the supervised PLS-DA analysis of targeted metabolomics under two conditions; (b) Heatmap overview of targeted metabolomics under two conditions.

Two-component systems (2CSTS) were confirmed to be involved in biofilm formation by transferring a phosphate from a conserved histidine to a conserved aspartate in the receiver domain of the response regulator^14^; which may account for the upregulation of L-aspartate and downregulation of L-histidine in our data (Figure 3a). Moreover, many differential metabolites are involved in the urea cycle and branches. It was reported that the Δ*ciaR* mutant of *Streptococcus sanguinis* had reduced biofilm formation by promoting the expression of arginine biosynthetic genes, especially the *argB* gene^15^. In addition, mutations in the genes encoding arginine decarboxylase (SpeA) and ornithine decarboxylase (SpeC) resulted in the loss of biofilm formation in *Yersinia pestis*, and the lack of citrullination in a peptidylarginine deiminase (PPAD) deletion mutant in *Porphyromonas gingivalis* enhanced biofilm formation, which may account for why arginine was remarkably increased while L-ornithine was decreased during biofilm formation in our study^16, 17^. We also found that polyamines such as arginine derivatives, including spermidine and putrescine, are required for biofilm formation^17–19^; this result is completely agreeable with the upregulation of these two metabolites during biofilm formation in our study (Figure 3b).

**Figure 3.**
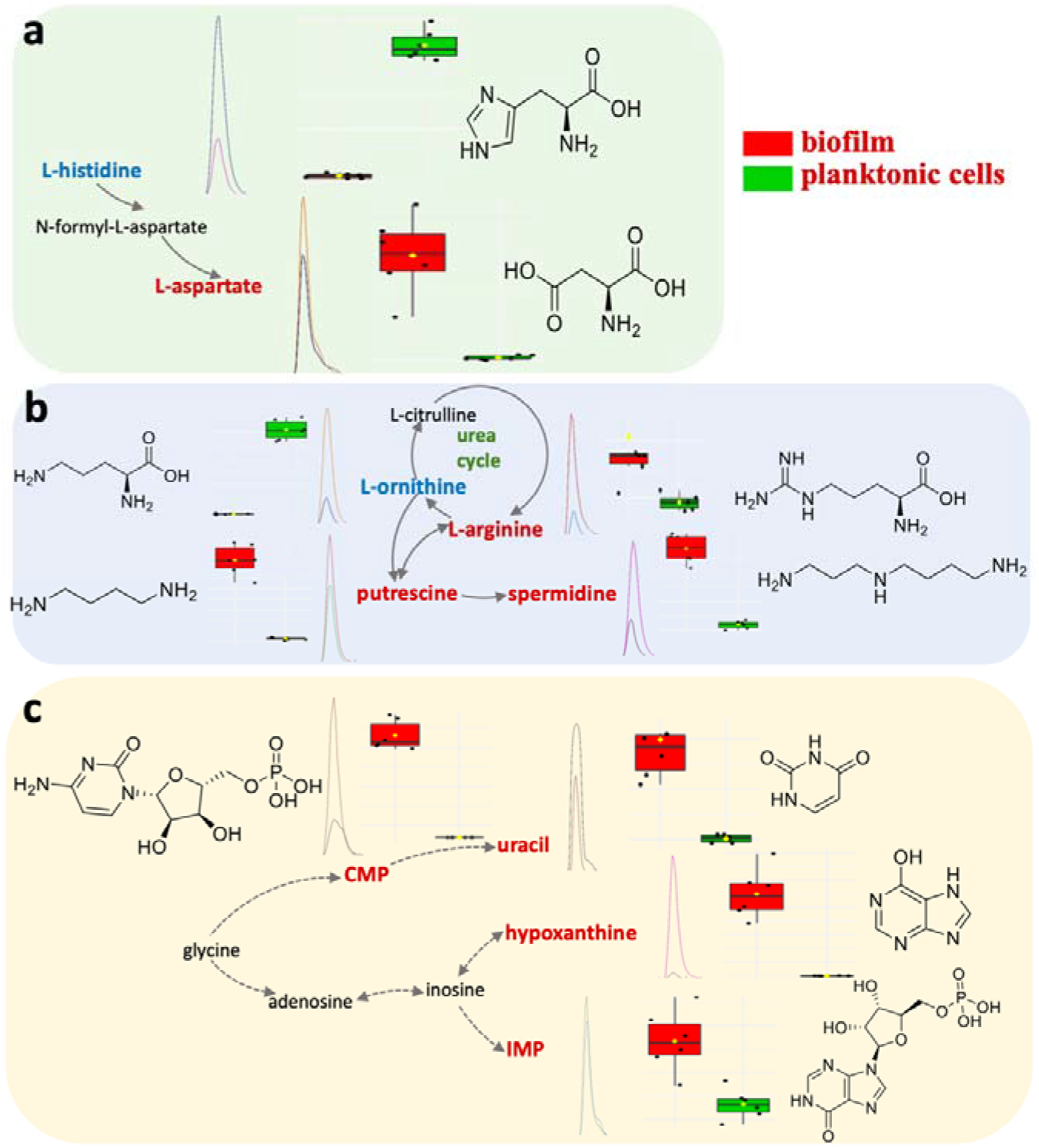
Mass spectrometry-based qualitative and quantitative characterization of key functional metabolites and associated metabolic pathways, whose levels were noticeably changed during biofilm formation. (a) Histidine metabolism; (b) urea cycle; (c) nucleotide biosynthesis.

Interestingly, most nucleotides and nucleosides, including CMP, IMP, uracil, AMP and hypoxanthine, were upregulated in biofilms; this result might indicate that biofilm formation requires more substrates to form EPS to survive under stressful environments (Figure 3c). In summary, using this new precision-targeted metabolomics method, our data illustrate unique metabolic characteristics of biofilm formation, and many key metabolites were confirmed to be directly or indirectly involved in biofilm formation. Given that metabolic reprogramming has been shown to be closely associated with biofilms, the biosynthesis of many metabolites might be the key effector for the formation process.

### Iron regulates biofilm formation in a concentration-dependent manner

Some studies have demonstrated that many metals exert substantial influences on biofilm formation ^20, 21^. Iron, a key metal for diverse living organisms, is an essential nutrient and cofactor for key proteins participating in critical biological processes during microbial growth ^22, 23^, and iron has been shown to be involved in regulating biofilm formation^24, 25^ in different bacterial cells.

To investigate the phenotypic features of the regulation of biofilm formation by iron, we employed different analytical methods to visualize biofilm formation in the presence of different levels of iron. Our data exactly demonstrated that biofilm formation was regulated by iron, and this effect was obviously concentration-dependent (Figure 4). Phenotype visualization methods (TEM, SEM, staining and CFU assay) were used to globally investigate the structure and physiological features of biofilms in the presence of different concentrations of iron. The treatments with different concentrations of iron were compared, and TEM and SEM images revealed that treatment with 1000 μM iron can significantly eliminate the fiber layer compared to the original culture medium. This result suggests that iron indeed regulates biofilm formation, which could be characterized by distinct changes in the biofilm structure (Figure 5). In addition, the staining and CFU assay confirmed that the capacity of biofilm formation was enhanced when the treated concentration of iron varied from 0 to 10 μM; however, the capacity of biofilm formation was inhibited with iron treatment at concentrations of 10 to 1000 μM. Excitingly, biofilm formation was completely inhibited with in the presence of 2000 μM iron. This finding showed was completely inhibited with in the presence of 2000 μ that iron beneficially regulates biofilm formation at a defined concentration (0 to 10 μM). Beyond this, overloading iron inversely regulates biofilm formation by inhibiting bacterial growth ^22, 23^ (Figure S3). Ideally, our finding is almost consistent with a previous study based on another opportunistic pathogen, *Pseudomonas aeruginosa*, allowing us to draw a short conclusion that a high concentration of iron could cause the perturbation of biofilm aggregates and escape of planktonic cells through oxidative damage^26^. In short, biofilm formation can be promoted by a defined concentration of iron but perturbed by a high concentration of iron (beyond 1000 μM); at a high concentration, iron inhibits bacterial growth by destroying the organized ultrastructure.

**Figure 4.**
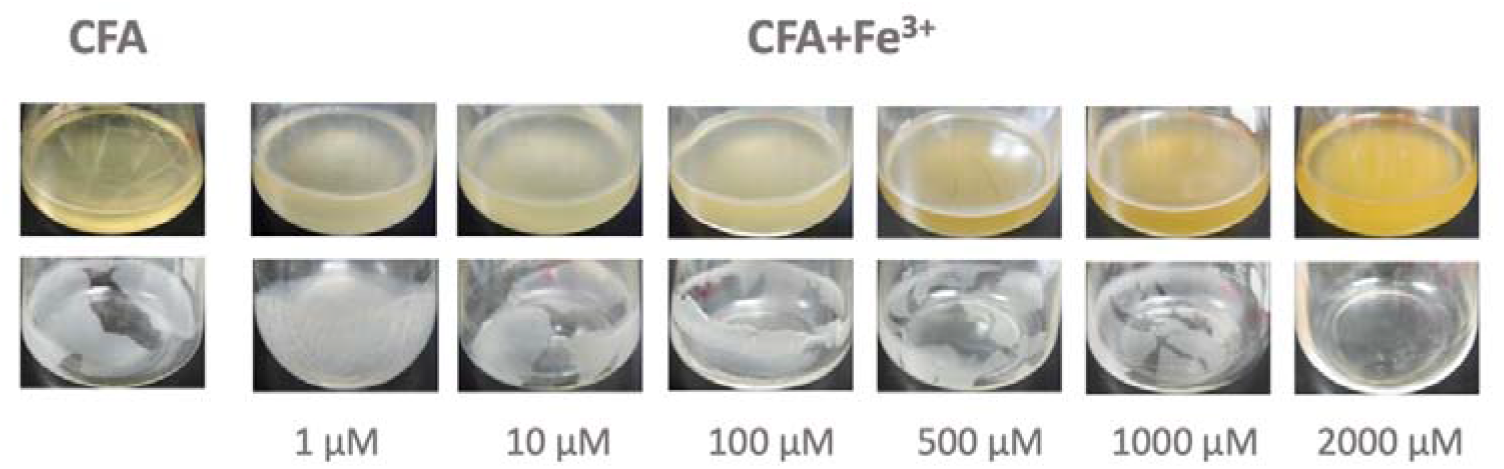
Phenotypic image that demonstrated that iron can inhibit biofilm formation in an obvious concentration-dependent manner, as characterized by the decrease in the observable layer of biofilm by increasing the concentration of iron supplemented in the culture medium. The key iron concentration to terminate biofilm formation in this culture system was 100 μM.

**Figure 5.**
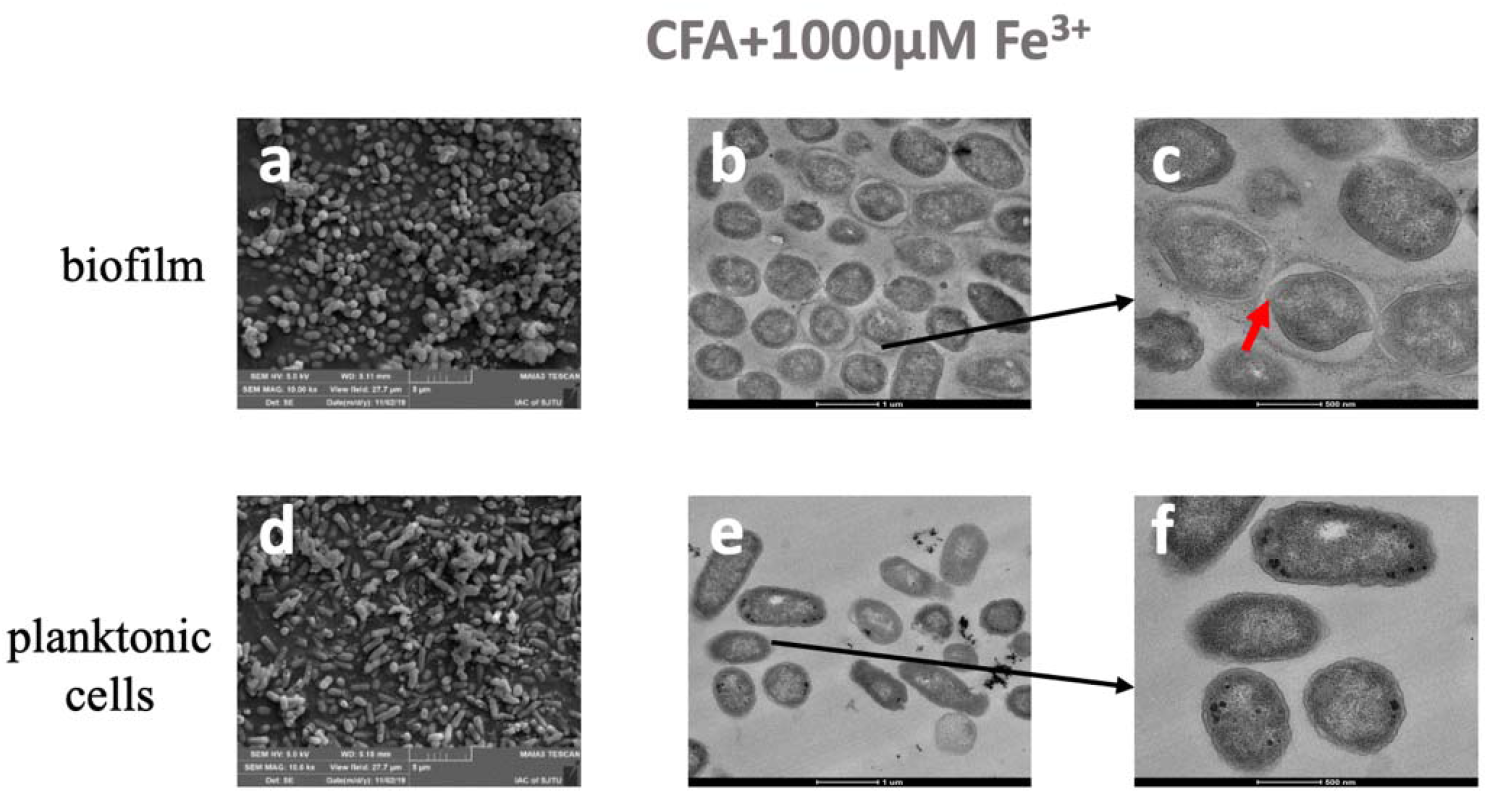
Comparison of phenotypic images revealed the distinct structure of biofilm pre- and posttreatment with 1000 μM iron. (a) SEM image of biofilm. (b and c) TEM image of biofilm; (d) SEM image of planktonic cells; (e and f) TEM image of planktonic cells.

### Iron regulates biofilm formation by targeting metabolic reprogramming

To elucidate the metabolic mechanism of iron regulation on biofilm formation, we measured the metabolites present during biofilm formation under treatments with different concentrations of iron. The metabolomics data indicate that the metabolic characteristics of biofilms were markedly modified by iron, and the modified pattern was obviously dependent on the iron concentration, which is almost consistent with the growth state of bacterial cells (Figure 6). A total of 23 functional metabolites with expressional-level changes mostly due to the metabolic regulation of iron on biofilm formation were identified (Table S3). As expected, the metabolic characteristics of biofilm formation were verified again by analyzing the functional metabolites that changed in the presence of iron, including L-isoleucine, L-leucine, CMP, L-tryptophan, L-histidine, L-aspartate, 5’-deoxy-5’-methylthioadenosine (5’-MTA), putrescine and spermidine. The changes in these functional metabolites were consistent with the phenotypes of biofilms, as the metabolite clusters of biofilms were significantly distinguished from those of planktonic cells with the initial increase in iron concentration. The metabolic pattern of the biofilm was preferably restored to that of the planktonic populations in the presence of a high concentration of iron. The results confirmed that iron modulates bacterial metabolism in a concentration-dependent manner by alternating the growth state of bacterial cells at different concentrations^13^. Our data further reveal that the iron-regulated biosynthesis of amino acids might be mostly involved in biofilm formation. For example, histidine has been reported to have the capacity to bind metal iron in bacterial cells ^27^. In addition, histidine kinase and urocanase are involved in histidine metabolism, which plays a crucial role in biofilm formation ^28, 29^, and histidine-functionalized silver nanoparticles could even eradicate biofilms of *Klebsiella pneumoniae*^30^. Moreover, leucine, which has been recorded as a component of biofilms ^31^, was incorporated into protein biosynthesis, and *E. coli* biofilms might use the localization of leucine to adapt to the static environment during its early stage development^32^. In addition, leucine aminopeptidase (LAP) enzymatic activity was inhibited by ion chelators^33^ (Figure 7a). Interestingly, polyamines such as spermidine and putrescine are not closely associated with biofilm formation; they also have the capacity to interact with iron^34^, protecting an organism’s DNA from the damage of reactive oxygen species (ROS) triggered by different agents, such as iron^35^, supporting our finding that polyamine plays a role in biofilm formation by interacting with iron (Figure 7b). We also noticed that the level of nucleotides and derivatives, including CMP and 5’-MTA, changed regularly in response to different iron concentrations, suggesting that iron might alternate biofilm formation by regulating the biosynthesis of critical materials (Figure 7c).

**Figure 6.**
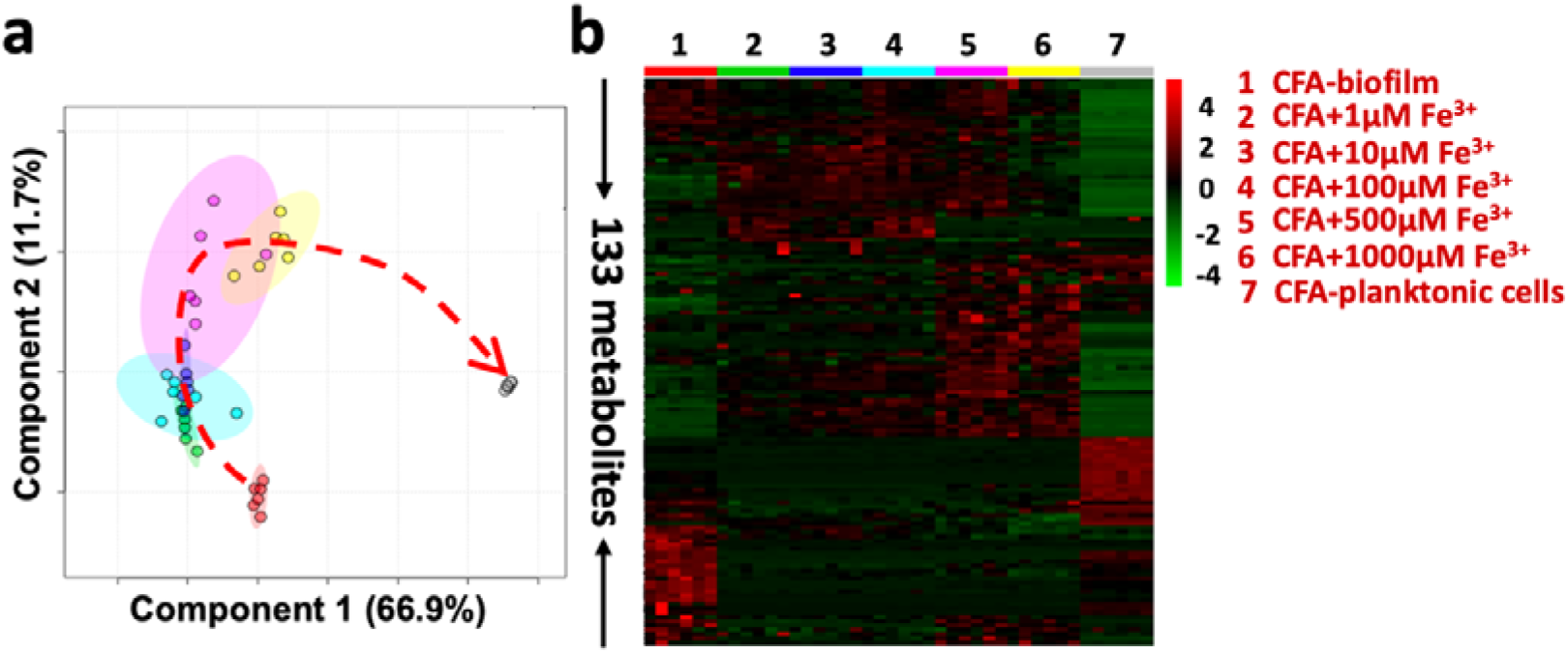
A targeted-metabolomics assay revealed that iron has the capacity to regulate the small-molecule metabolism of biofilms in an iron concentration-dependent manner. (**a)** Score plot resulting from the supervised PLS-DA analysis of precision-targeted metabolomics; (**b)** heatmap-overview of metabolomes.

**Figure 7.**
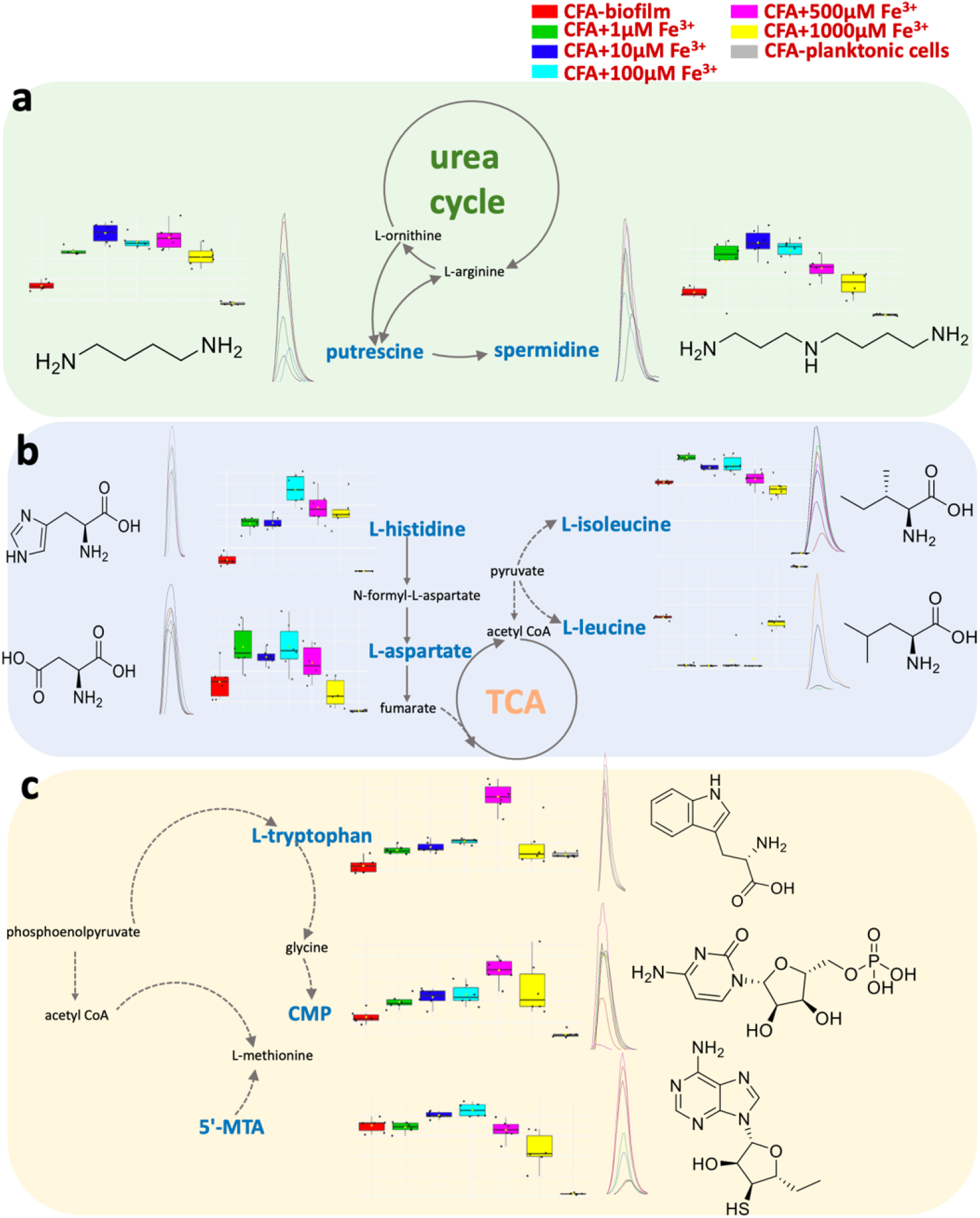
Mass spectrometry-based qualitative and quantitative characterization of key functional metabolites and associated metabolic pathways, whose levels were noticeably changed during biofilm formation in the presence of different concentrations of iron. (a) Urea cycle; (b) TCA cycle; (c) nucleotide biosynthesis.

### Bioavailable iron is transported into bacterial cells by siderophores to regulate biofilm formation

Siderophores are chemically diverse metabolites with high affinity for iron and are used by microorganisms as an iron acquisition system to regulate the bioavailability of Fe^3+^ from the iron-limited environment^36–38^. Because bacterial cells primarily take up iron in a siderophore-dependent pathway, we hypothesized that iron is first pumped into bacterial cells by the siderophore transport system and that iron can regulate the biosynthesis of functional metabolites that are key players in biofilm formation. To verify our hypothesis, we employed an LC-QTOF MS method to analyze four key siderophores biosynthesized by the UTI89 strain, including yersiniabactin, linear enterobactin, di-glucosylated-salmochelin and HPTT-COOH, which have a robust capacity to chelate iron ^12, 39–41^ (Figure S4 **and** Figure 8). Our data revealed that four siderophores significantly interacted with iron; the levels of yersiniabactin, di-glucosylated-salmochelin and HPTT-COOH were largely decreased with increasing iron concentration, and the level of linear enterobactin was clearly increased by iron (Figure 8). The results suggested that the consumption of siderophores can increase the bioavailability of iron to bacterial cells and that the iron in cells can resume its role in regulating the biosynthesis of functional metabolites *in vivo.* This finding is almost consistent with a previous publication, which indicated that a standardized range of iron supplementation can modulate *E. coli* iron scarcity responses^13^. In addition, considering the dynamic interactions between iron and the four siderophores, at the initial stage, bacterial cells prefer to utilize nonconserved siderophores (yersiniabactin, di-glucosylated-salmochelin and HPTT-COOH) that can be produced by pathogenic *E. coli* strains to regulate iron intake. While the majority of nonconserved siderophores are completely consumed, bacterial cells are supposed to recruit conserved enterobactin, which can be produced by both pathogenic and nonpathogenic strains, to chelate iron. In short, such a mechanism allows bacterial cells to maintain the capability to utilize iron, which is essential to maintain survival in a variety of environments (Figure S3). Indeed, our data again show that microorganisms mostly adopt siderophore-dependent pathways to utilize iron from different growth environments ^42, 43^.

**Figure 8.**
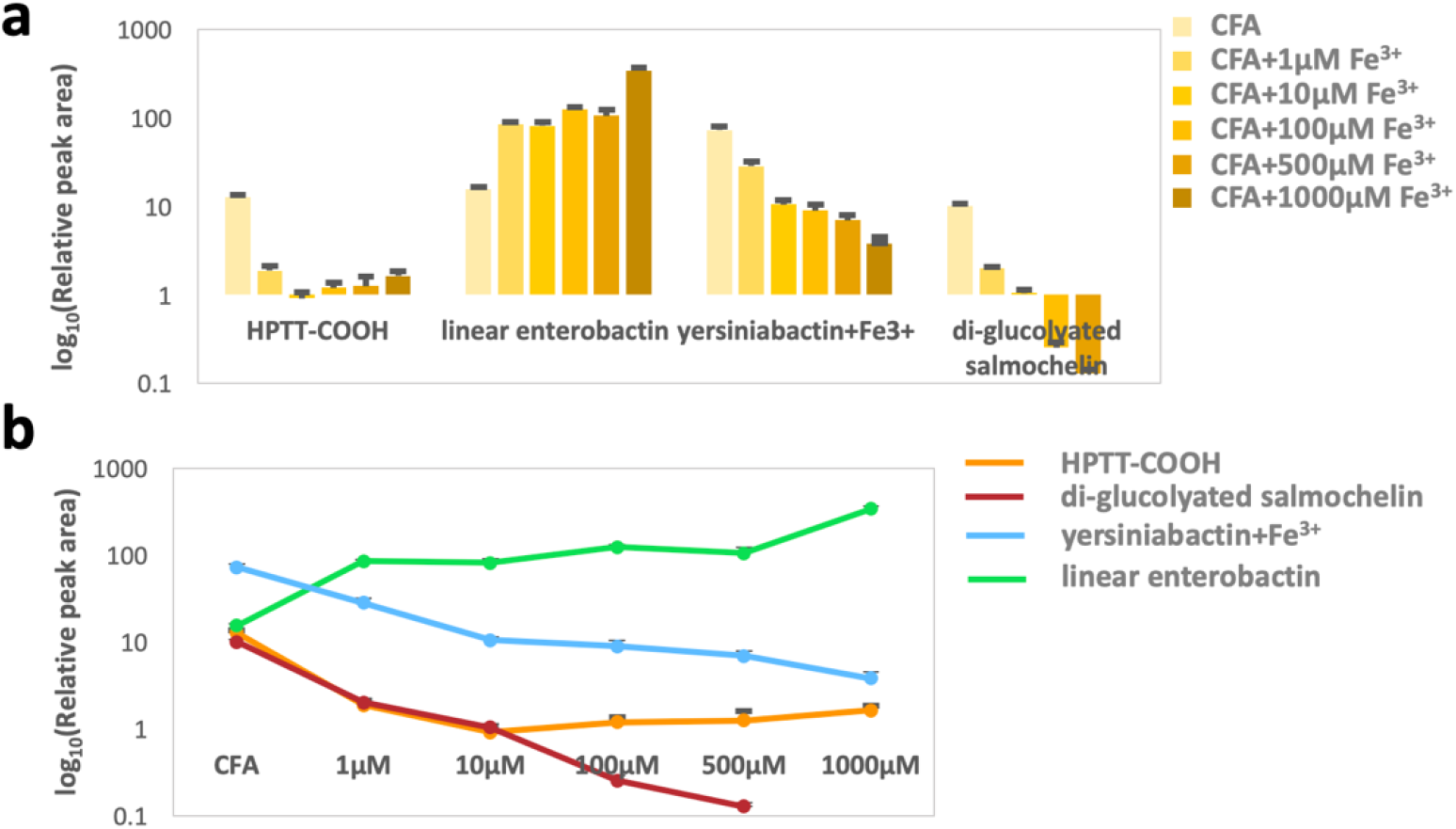
Expressional-level changes of the selected siderophores are regulated by the iron in a concentration-dependent manner. (a) Bar plot; (b) line plot.

Since iron has been observed to significantly regulate the biosynthesis and expression of functional metabolites that mostly contribute to the unique metabolic characteristics of biofilm formation and iron must be transported into bacterial cells by siderophores before resuming such a role, we wanted to learn whether siderophores, as secondary metabolites biosynthesized by primary metabolites, have the capacity to interact with the key functional metabolites of biofilm formation. Thus, we examined the correlation between the expression levels of nine functional metabolites and four siderophores. The results show that the biosynthesis of siderophores is closely associated with different functional metabolites (Figure 9), which is consistent with our previous study ^44^. The data showed that linear enterobactin had a negative correlation with most of the nine functional metabolites compared to the other siderophores (yersiniabactin, di-glucosylated-salmochelin and HPTT-COOH), which might be explained by the role of linear enterobactin in transporting iron into bacterial cells in the late stage of biofilm formation. Altogether, the siderophore-metabolite correlation analysis demonstrates that siderophores, chemically diverse secondary metabolites, can transport iron into bacterial cells, and siderophores are thought to be further involved in biofilm formation by regulating the interaction of functional metabolites with iron (Figure 10).

**Figure 9.**
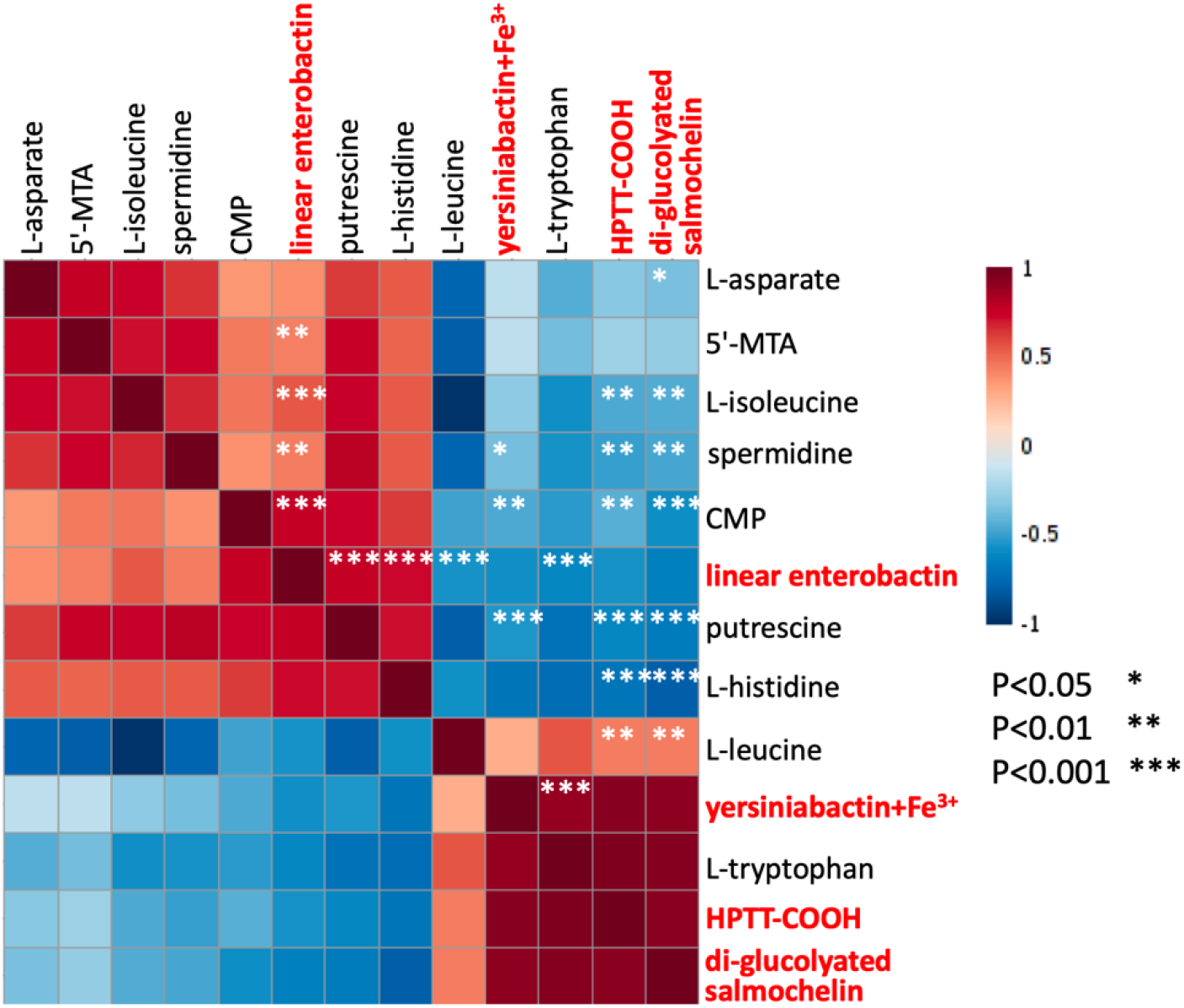
Expressional correlation analysis with the interactive heat map verified that the biosynthesis of siderophores regulates the unique pattern of functional metabolites, and iron might be recruited by cells as an essential intermediate to engage in such interactions.

**Figure 10.**
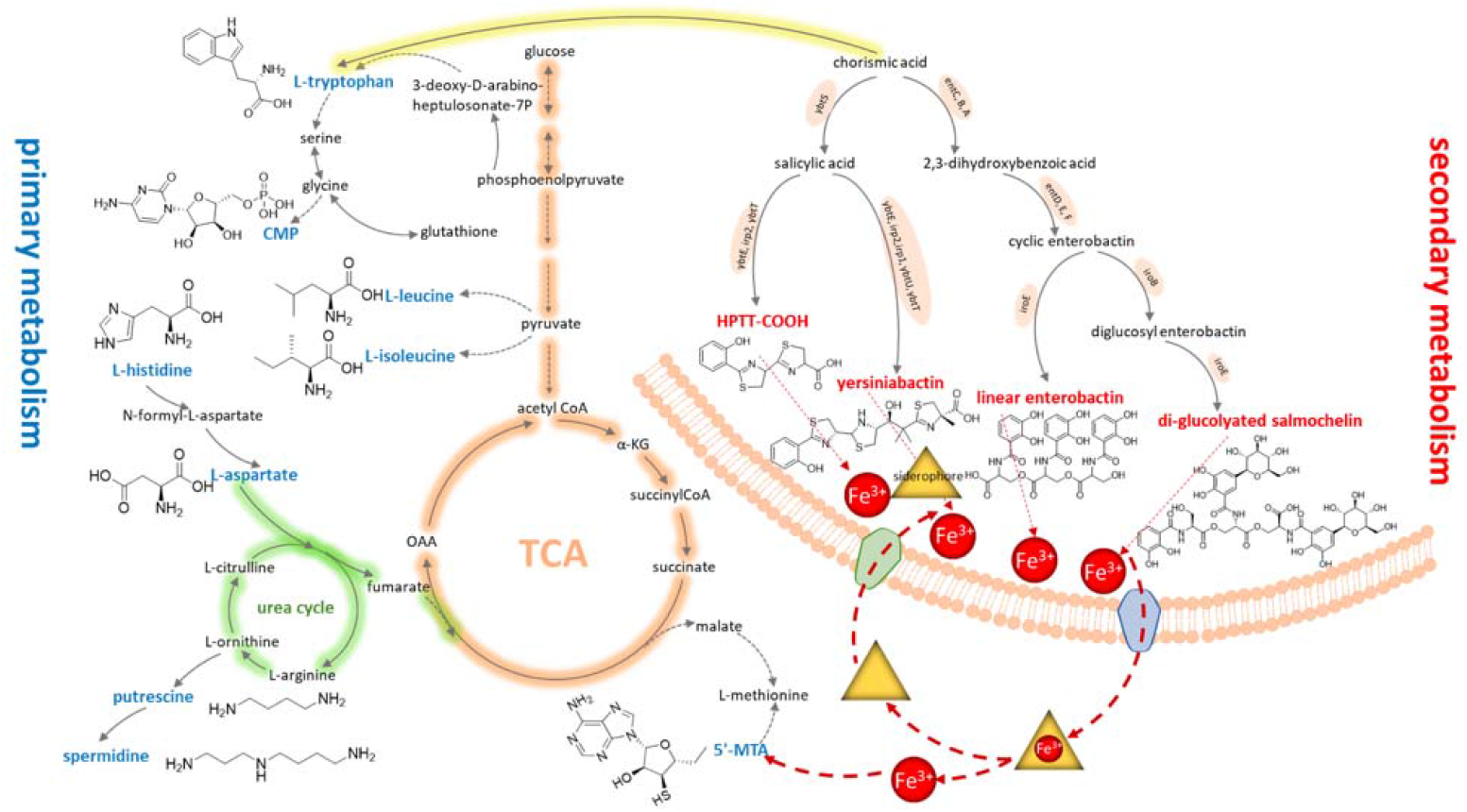
Bioavailable iron is primarily transported by siderophores into bacterial cells to further direct the biosynthesis and expression of functional metabolites, which eventually regulate biofilm formation.

### Iron mediated the biosynthesis and expression of functional metabolites to regulate biofilm formation

Nine functional metabolites have been shown to be involved in biofilm formation, and these metabolites have been observed to interact with iron utilized by the siderophore-dependent pathway. To elucidate the regulatory role of functional metabolites in biofilm formation, we supplemented the culture medium with more functional metabolites to observe the influence of the metabolites on biofilm formation. Surprisingly, four functional metabolites (L-tryptophan, 5’-MTA, spermidine and CMP) exerted inhibitory effects on biofilm formation, whereas L-leucine could promote biofilm formation (Figure 11). Generally, L-tryptophan has been reported to prevent *E. coli* biofilm formation via catalysis into indole ^45, 46^, and exogenous spermidine can reduce biofilm formation in a NspS/MbaA-dependent manner in *Vibrio cholerae*^47, 48^. Excitingly, our finding is the first to reveal the direct regulatory roles of 5’-MTA, CMP and L-leucine in biofilm formation. However, 5’-MTA was found to alter the quorum sensing (QS) signal system, which might further modulate biofilm formation and virulence in several bacterial strains ^49, 50^. Leucine was also shown to accumulate in the early stage of *E. coli*-related biofilm formation, and the leucine-protein complex has been applied to measure the biomass production of biofilms ^30, 31^. In short, we were able to verify the regulatory roles of five functional metabolites in biofilm formation, and these roles likely occur due to the role of iron treatment in regulating the biosynthesis and expression of functional metabolites during biofilm formation. We will further explore the molecular mechanisms of the regulation of biofilm formation by functional metabolites in our future work. Nevertheless, L-tryptophan, 5’-MTA, spermidine, CMP and L-leucine, in the present study, were shown to directly regulate biofilm formation by interacting with iron utilized by the siderophore-dependent pathway; the biosynthesis of these molecules can be targeted to design and develop novel inhibitors with the innovative potential to answer biofilm-related questions in different fields.

**Figure 11.**
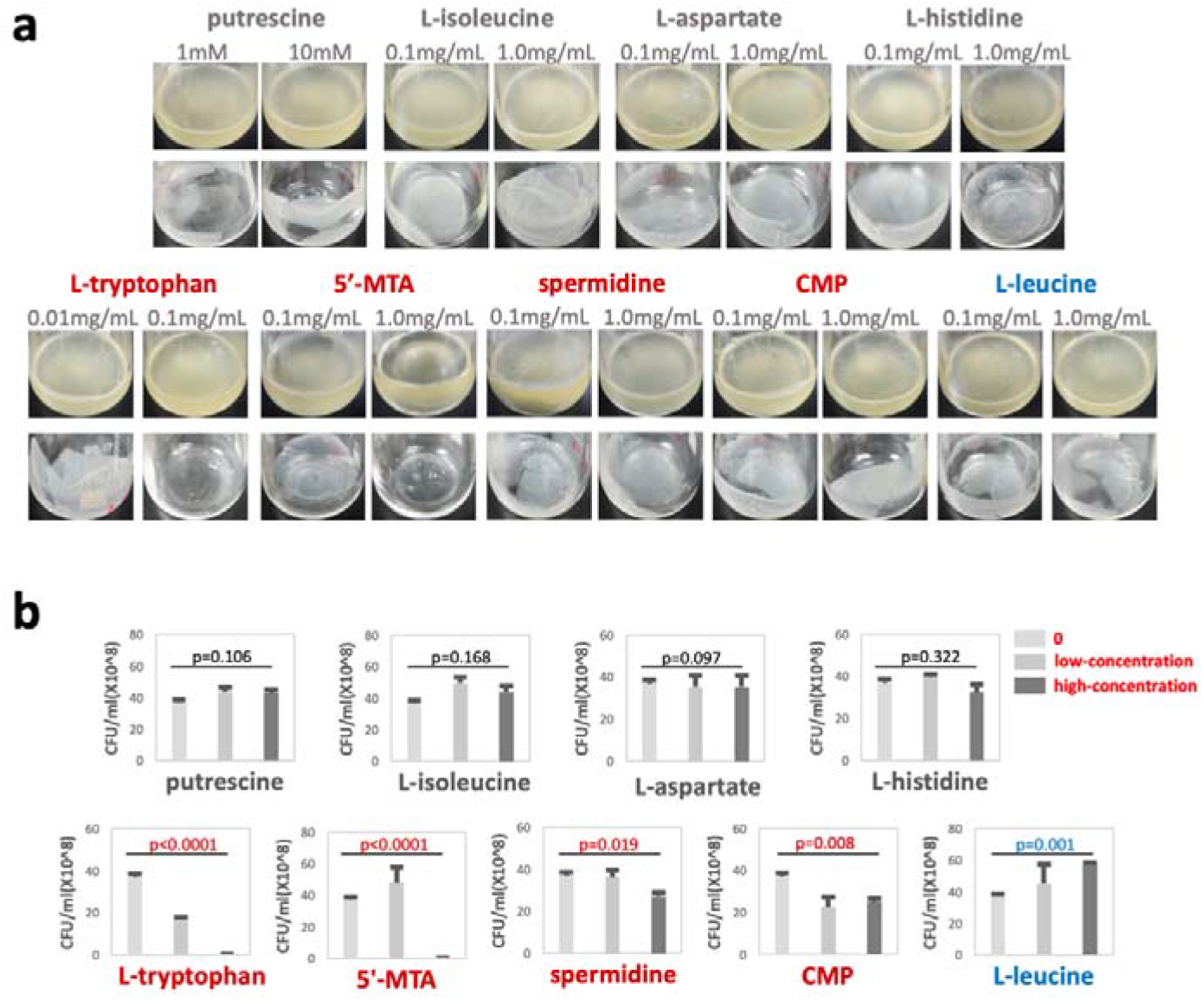
Five functional metabolites were shown to directly regulate biofilm formation in different patterns. L-Tryptophan, 5’-MTA, spermidine and CMP were experimentally validated to inhibit biofilm formation, which can be terminated with increasing treatment concentrations of the four functional metabolites. Another functional metabolite (L-leucine) was observed to promote biofilm formation. (A) Image of the phenotype; (B) CFU determination.

## Discussion

Over the past decades, biochemists in different niches have made great efforts in biofilm formation research, allowing us to comprehend the potential mechanism of biofilm formation ^1, 2, 6^. However, because biochemical complexity occurs with biofilm formation, the mechanism of biofilm formation in different microorganisms has not been completely elucidated. Consequently, biofilm formation causes challenging problems largely present in different areas; for example, environmental pollution, infectious diseases, agricultural production, industrial contamination and antibiotic resistance ^3–5^. To overcome the difficult challenges associated with biofilm formation, we sought to decipher the mechanism of biofilm formation from a metabolic perspective because our recent study confirmed that substantial metabolic reprogramming often occurs during biofilm formation^6^. Following this new discovery, our study aimed to further decipher the potential mechanism of metabolic regulation during biofilm formation to enable the development of a novel strategy that regulates functional metabolites to terminate biofilm formation to tackle the challenging problems mentioned above. Imaging methods facilitated phenotype visualization to confirm that biofilm formation was stimulated by the bacterial strain (UTI89 *E. coli*) cultivated in the optimized culture medium, and we efficiently visualized that biofilm formation allows bacterial cells to reorganize in a unique structure, differentiating these cells from planktonic cells ^6, 10^. Our new targeted metabolomics method^11^ was used to precisely investigate the metabolic characteristics of biofilm formation under different conditions. Many metabolites and associated metabolic pathways were characterized accordingly, and these pathways were shown to be involved in biofilm formation. In particular, many metabolites could be defined as functional molecules that were shown to regulate biofilm formation; for example, L-histidine was remarkably decreased, and biofilm formation can be induced by the transfer of a phosphate from histidine to aspartate by 2CSTS ^14^. In our study, we found that the expression level of L-histidine in biofilms was quite lower than that in planktonic cells, which argued that biofilm formation often incurs the metabolic cost of L-histidine. In another observation, we noticed that the expression levels of polyamines (spermidine and putrescine) were significantly increased, which agrees with previous findings ^17, 18^, suggesting that polyamines might be functional molecules for phenotyping biofilm formation. Furthermore, our data reveal that biofilm formation is obviously affected by bioavailable iron in a concentration-dependent manner through the regulation of many functional metabolites; for example, biofilm formation is regulated by iron through binding sites and the DNA protection role of polyamines (spermidine and putrescine) ^34, 35^. To correlate siderophores with the bioavailability of iron, we found that iron is mostly transported into bacterial cells by a siderophore-dependent pathway. We are the first to verify that bacterial cells mostly utilize nonconserved siderophores (yersiniabactin, di-glucosylated-salmochelin and HPTT-COOH) to intake bioavailable iron in an iron-limited environment. Instead, bacterial cells switch to recruit conserved siderophores (linear enterobactin) after these nonconserved siderophores are depleted to undertake the role of transporting iron to maintain normal growth in an iron-rich environment. Given that biofilm formation involves the interactions of iron and functional metabolites, siderophore-functional metabolite correlation analysis was further performed to understand the roles of siderophores in biofilm formation. The phenotypic data show that siderophores considerably affect the expression of functional metabolites regulated by bioavailable iron, which is mostly underlined in biofilm formation. As expected, five iron-directed functional metabolites, including L-tryptophan, 5’-MTA, spermidine, CMP and L-leucine, were discovered to regulate biofilm formation. These five metabolites can be regarded as novel metabolic effectors that mediate biofilm formation under the regulation of bioavailable iron (1-10 μM) and maintain the health of microorganisms ^51–54^. Consequently, increasing the exposure level of iron (over 100 μM) to dysregulate the biosynthesis and expression of these functional metabolites and terminate biofilm formation should be substantially beneficial to the broad fields in the life sciences by allowing all the challenging problems related to biofilm formation to be tackled; for example, infections frequently occur in the human community involving UTIs and oral candidosis7, ^55^. Taken together, we are the first to elucidate the mechanism of biofilm formation through siderophores, which enable iron to regulate functional metabolites, as iron is chelated and transported into bacterial cells. For the first instance, iron selectively directs the biosynthesis and expression of functional metabolites; consequently, functional metabolites are recruited by bacterial cells to regulate biofilm formation (Figure 12). Such discovery allows the further design and development of new strategies to inhibit biofilm formation by the regulating functional metabolites.

**Figure 12.**
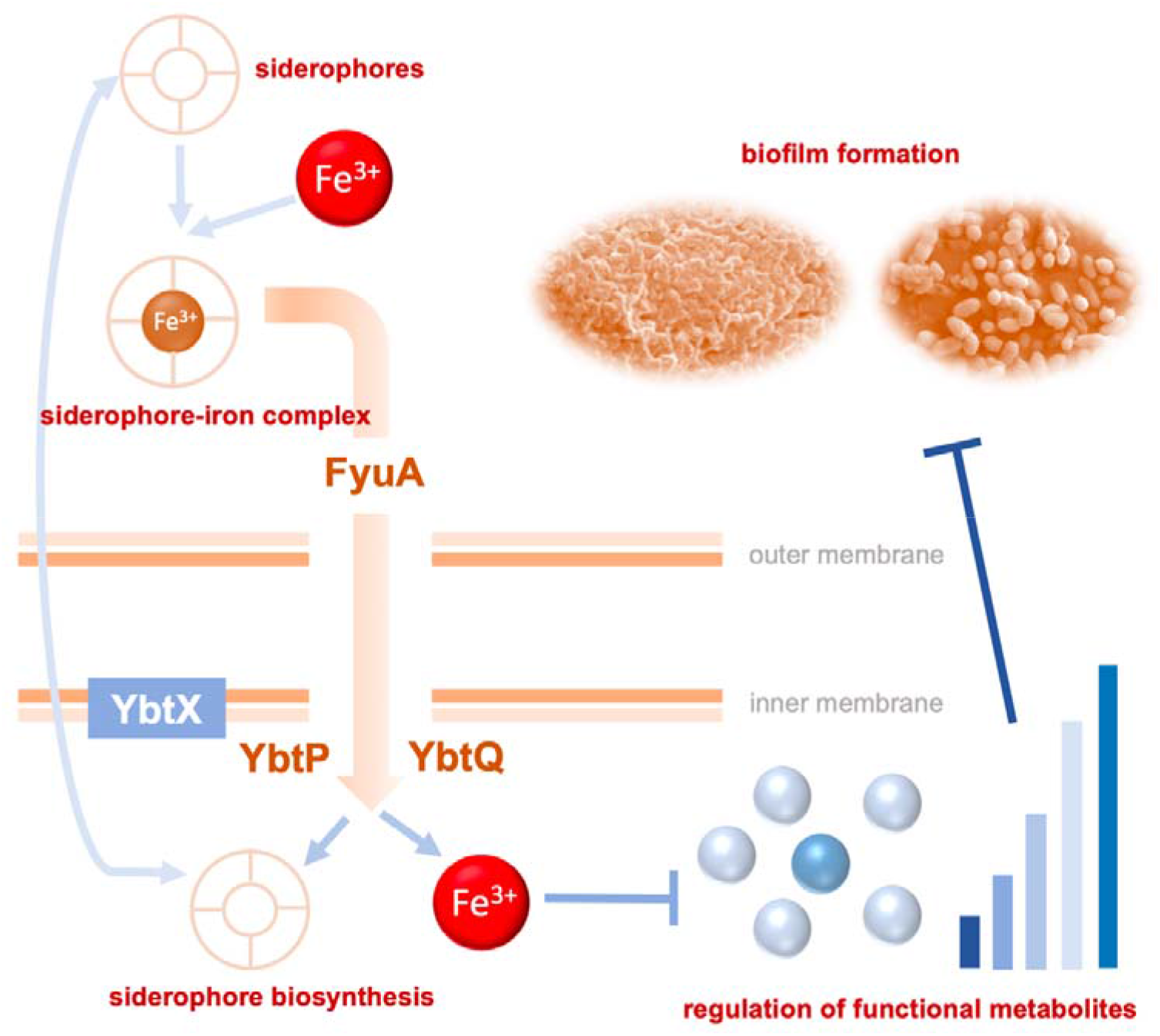
Schematic illustration of the biofilm formation mechanism in bacterial organisms. Briefly, the metal siderophores chelate and transport iron into bacterial cells. Then, iron is recruited to direct the biosynthesis and expression of functional metabolites, which subsequently regulate biofilm formation in a unique organized structure by dysregulating the metabolic characteristics of planktonic cells.

## Conclusion

Biofilms triggered by diverse microorganisms are still a severe challenge to the broad community because the formation mechanism has not been completely elucidated. However, biofilm formation often leads to the unexpected outbreak of numerous harmful problems, including environmental pollution, agricultural production, industrial contamination, and infectious diseases. In this study, we employed a functional metabolomics method to precisely identify five functional metabolites that can regulate biofilm formation under the regulation of bioavailable iron, which is mostly transported into bacterial cells through the siderophore-dependent pathway. In short, these innovative findings should provide novel insights into the mechanism of biofilm formation and can largely assist in the development of strategies to answer key questions regarding biofilms in different niches.

## Acknowledgments

This work was supported by the National Key R&D Program of China (No. 2017YFC1308600 and 2017YFC1308605), the National Natural Science Foundation of China Grants (No. 81274175 and 31670031), the Startup Funding for Specialized Professorship Provided by Shanghai Jiao Tong University (No. WF220441502).

---

**Figure S1.**
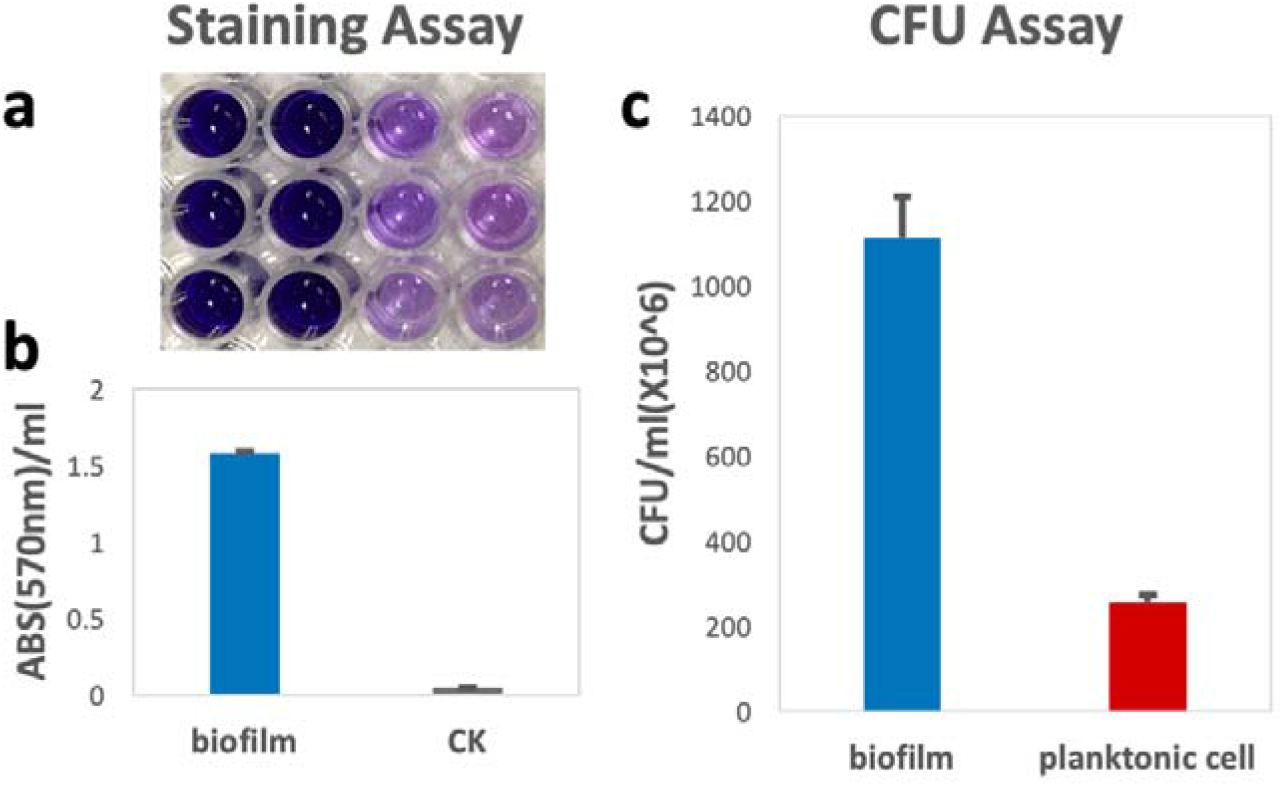
Staining combined with the CFU assay was adopted to quantitatively assess biofilm formation. (**a)** Crystal violet staining; **(b)** absolute value (OD570 nm) of crystal violet in (a) was quantitatively measured; (**c)** CFU assay.

**Figure S2.**
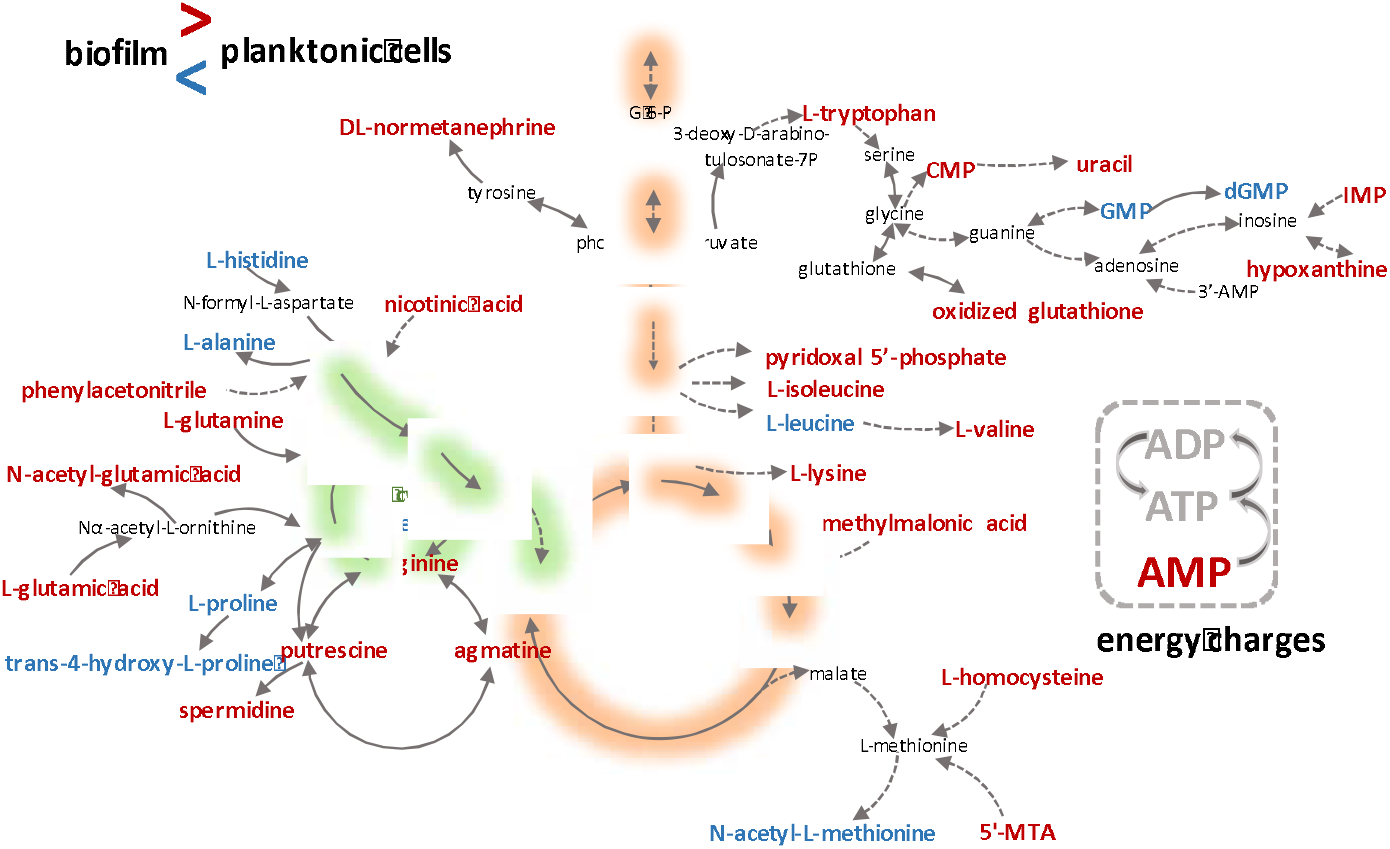
The most alternated metabolites and associated metabolic pathways were globally mapped and were observed to be involved in substantial metabolic reprogramming to permit biofilm formation.

**Figure S3.**
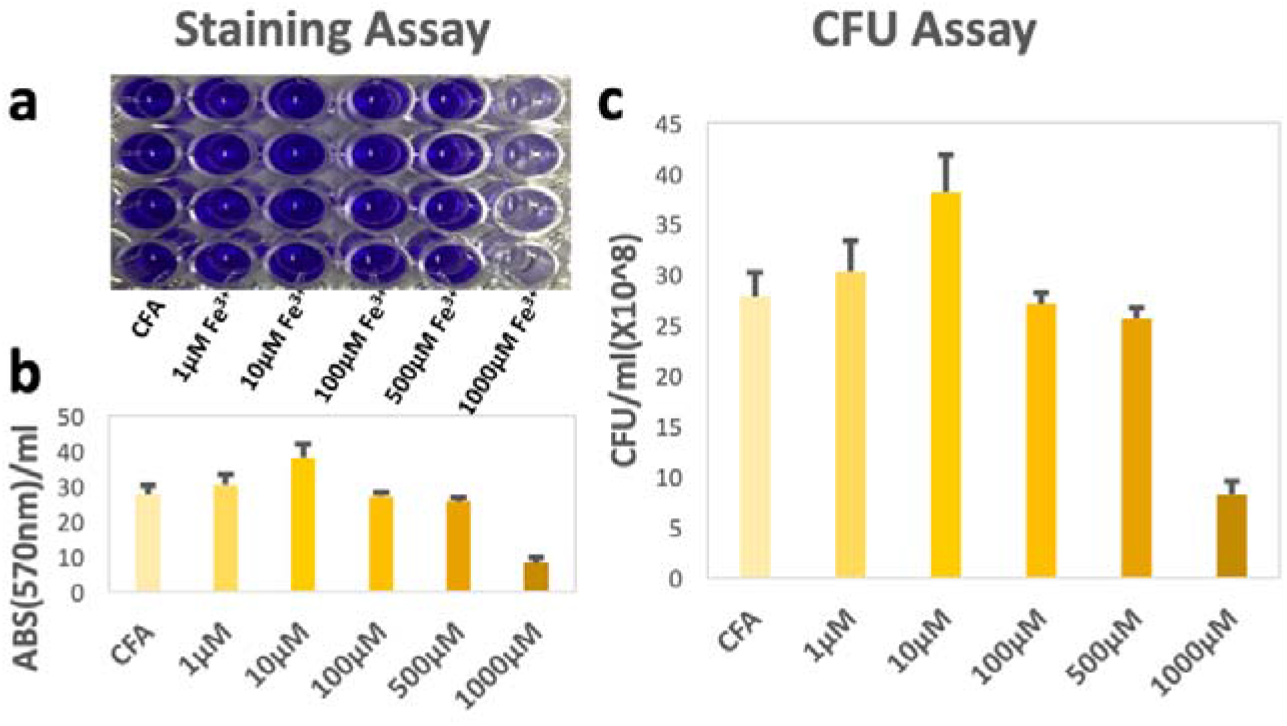
Staining combined with the CFU assay was applied to quantitatively assess biofilm formation under different conditions. (a) Crystal violet staining; (b) absolute value (OD570nm) of crystal violet in (a) was quantitatively measured; (c) CFU assay.

**Figure S4.**
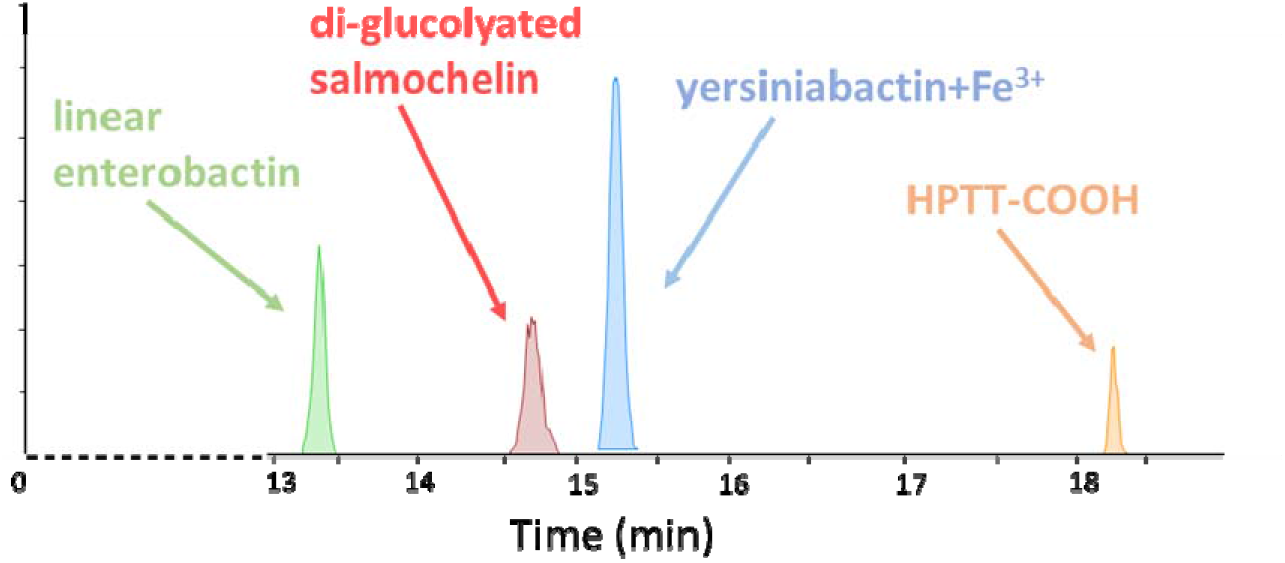
LC-QTOF MS profile method for the determination of four key siderophores

**Table S1.**
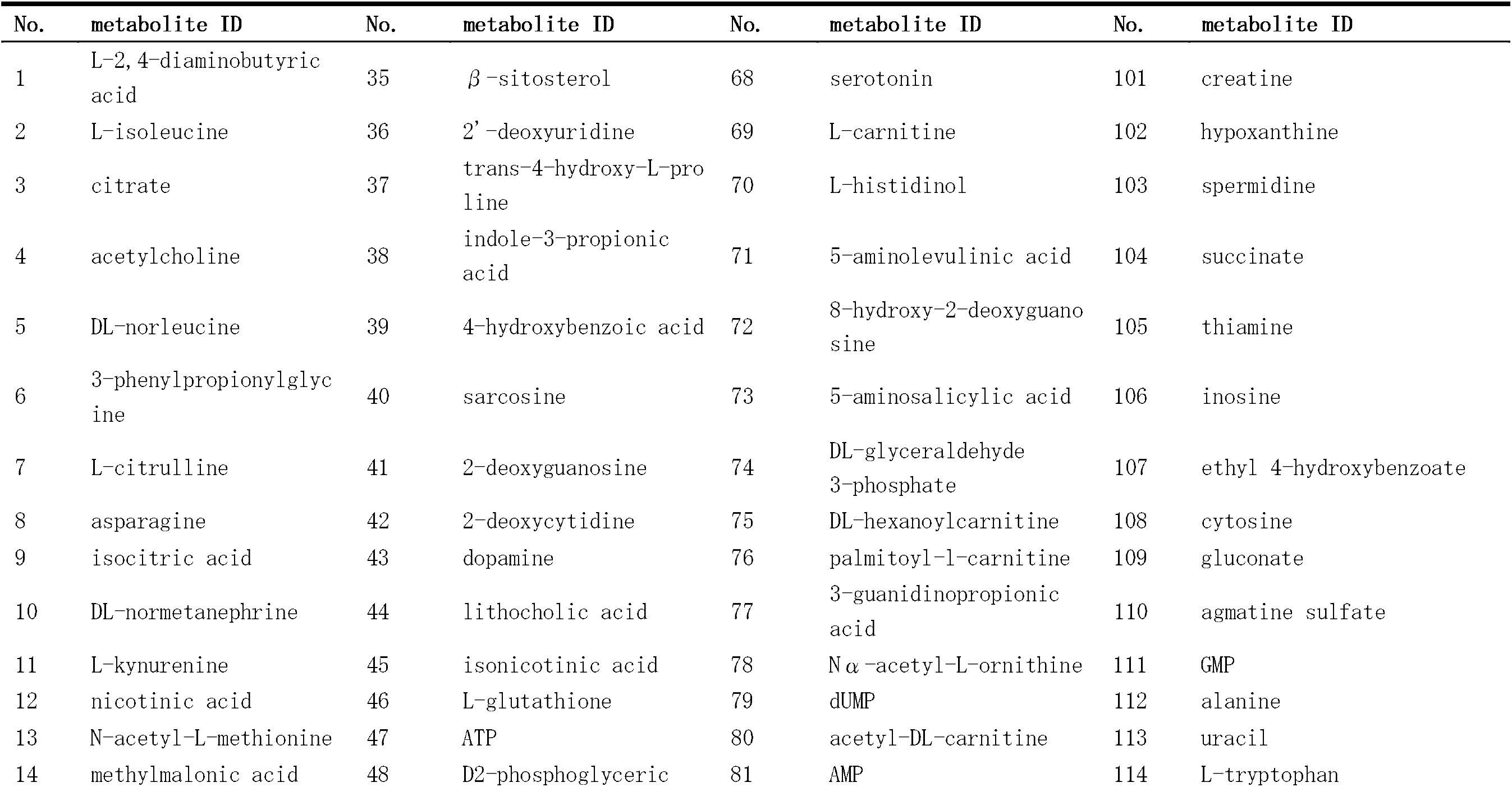
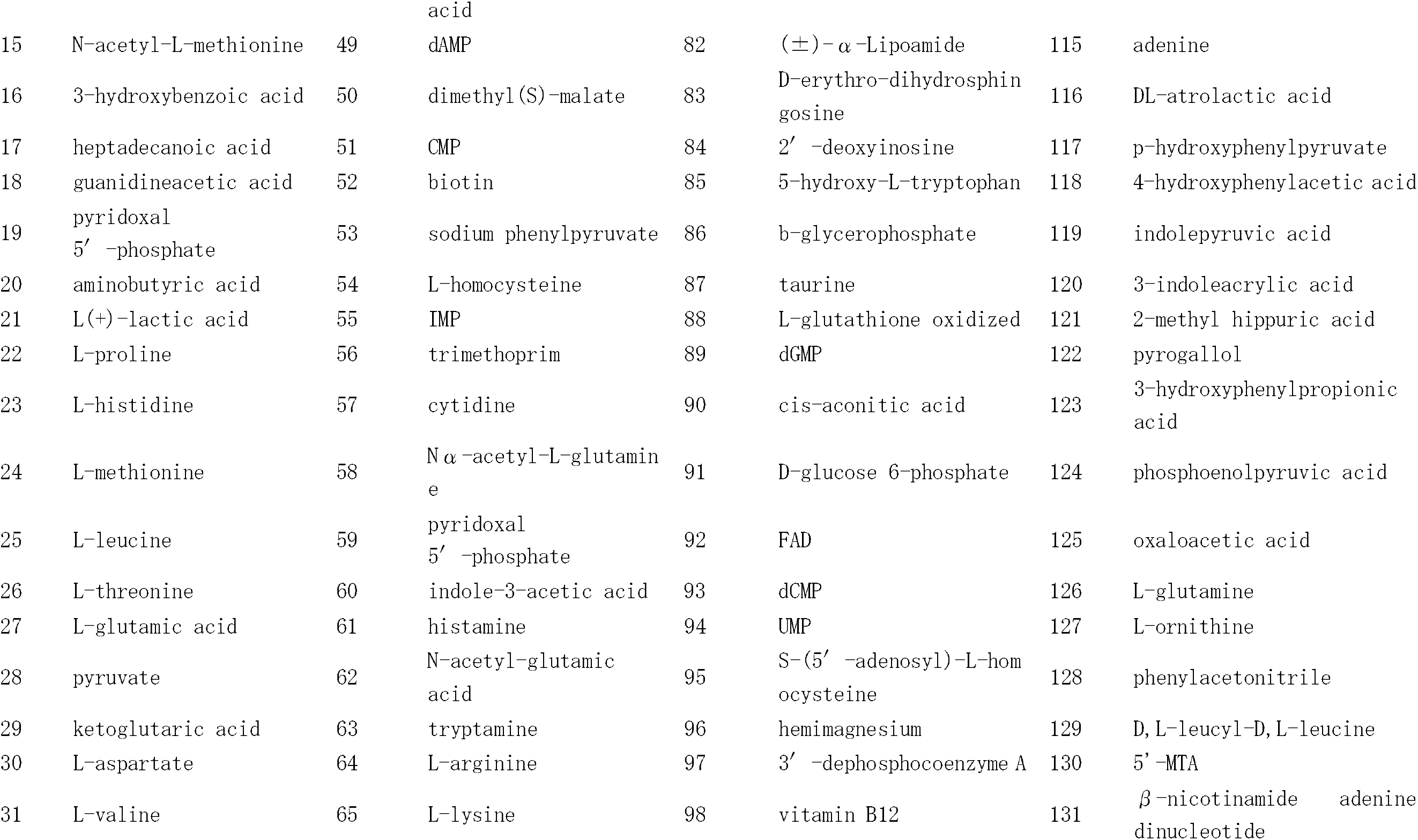
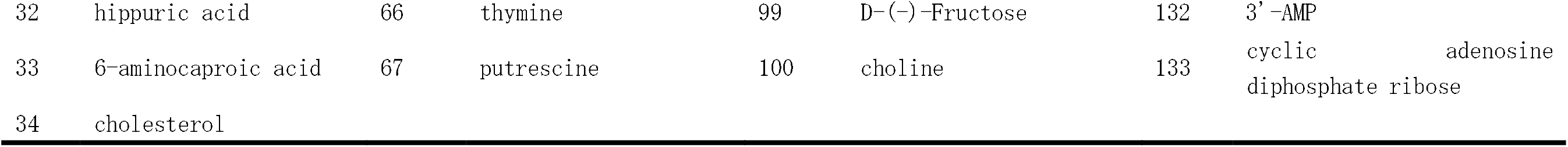
LC/TQ MS-based analysis of well-defined metabolites of interest used to investigate the metabolic characteristics of biofilm formation.

**Table S2.**
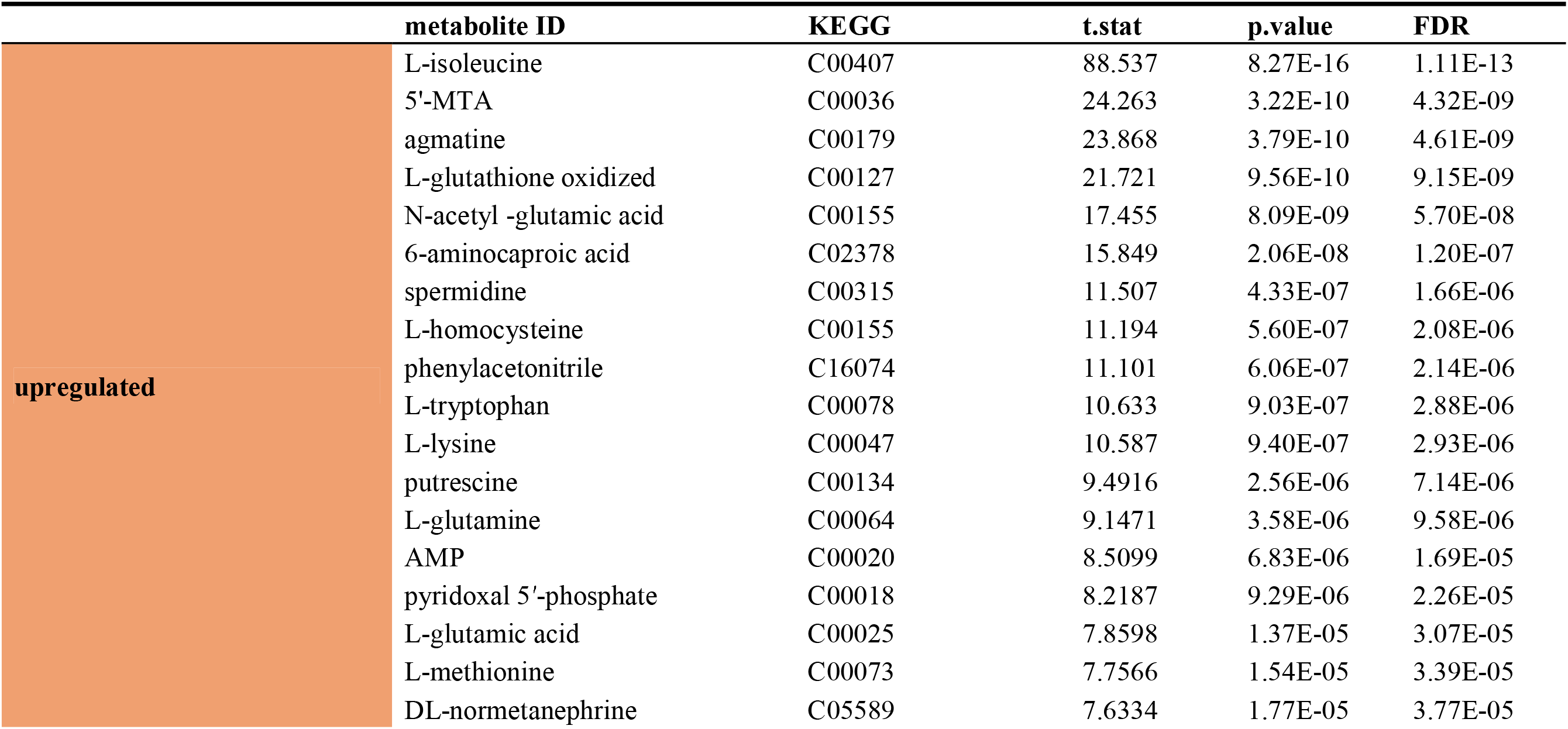
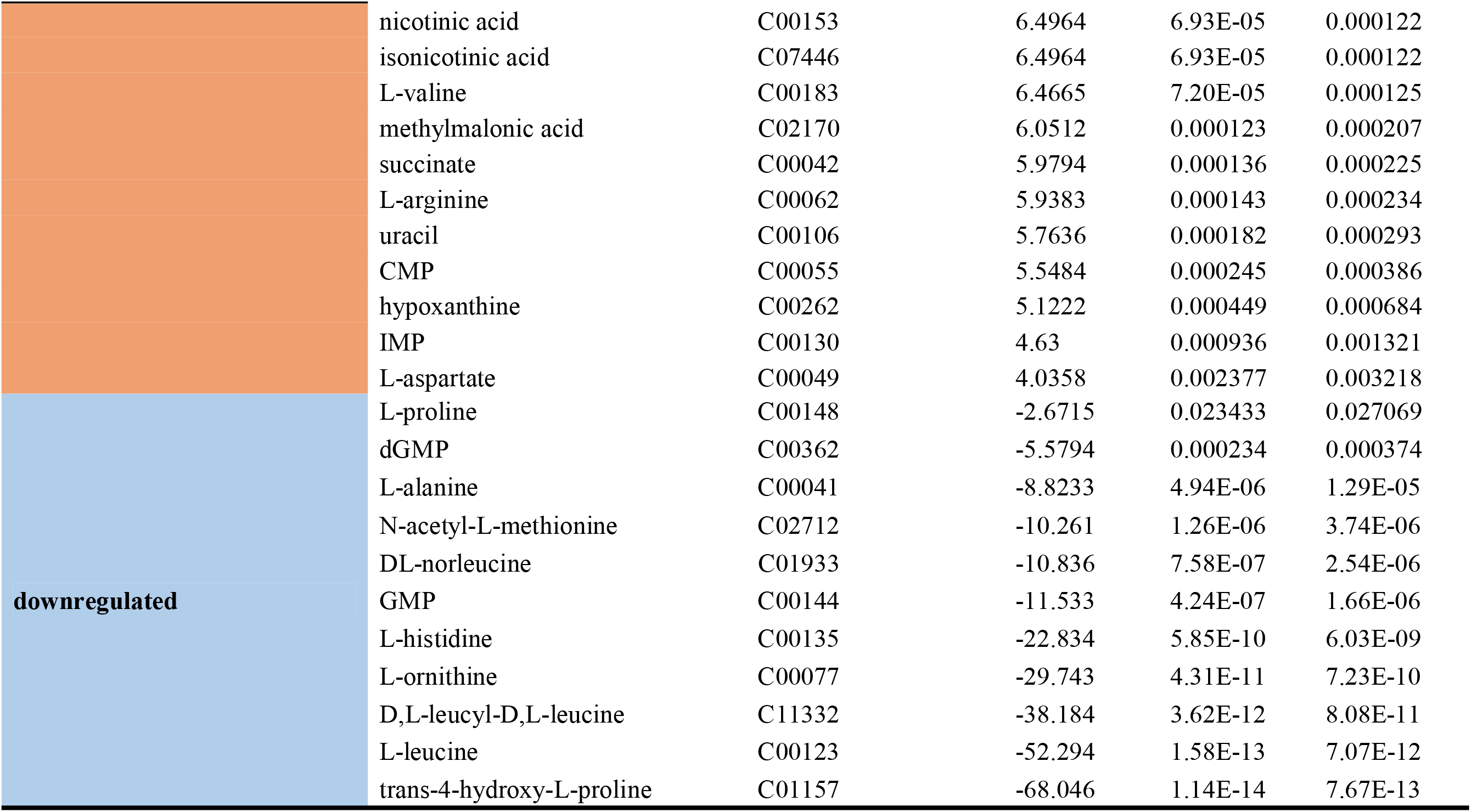
Differential metabolites confirmed using a targeted metabolomics method to distinguish biofilms from planktonic cells.

**Table S3.** Key metabolites regulated by iron and involved in the metabolic characteristics of biofilm formation.

|  | metabolite ID | KEGG | f.value | p.value | FDR |
| --- | --- | --- | --- | --- | --- |
| **biofilm formation-related** | L-isoleucine | C00407 | 161.42 | 5.27E-24 | 5.43E-23 |
|  | L-leucine | C00123 | 117.62 | 1.04E-21 | 8.23E-21 |
|  | L-aspartate | C00049 | 12.875 | 1.25E-07 | 1.76E-07 |
|  | CMP | C00055 | 14.534 | 3.01E-08 | 4.54E-08 |
|  | putrescine | C00134 | 66.51 | 1.13E-17 | 4.86E-17 |
|  | L-histidine | C00135 | 42.61 | 1.16E-14 | 3.30E-14 |
|  | spermidine | C00315 | 20.458 | 4.06E-10 | 7.45E-10 |
|  | L-tryptophan | C00078 | 140.46 | 5.43E-23 | 4.55E-22 |
|  | 5'-MTA | C00036 | 61.565 | 3.84E-17 | 1.43E-16 |
| **others** | citrate | C00158 | 35.381 | 1.87E-13 | 4.82E-13 |
|  | asparagine | C00152 | 45.881 | 3.75E-15 | 1.12E-14 |
|  | L-threonine | C00188 | 12.079 | 2.57E-07 | 3.46E-07 |
|  | 2'-deoxyuridine | C00460 | 549.9 | 4.13E-33 | 9.22E-32 |
|  | ATP | C00002 | 22.638 | 1.05E-10 | 2.10E-10 |
|  | D2-phosphoglyceric acid | C00197 | 20.965 | 2.93E-10 | 5.62E-10 |
|  | indole-3-acetic acid | C00954 | 16.793 | 5.14E-09 | 8.30E-09 |
|  | L-carnitine | C00134 | 21.966 | 1.57E-10 | 3.06E-10 |
|  | cis-aconitic acid | C00417 | 16.847 | 4.94E-09 | 8.17E-09 |
|  | FAD | C00318 | 24.742 | 3.11E-11 | 6.62E-11 |
|  | D-(-)-fructose | C00095 | 108.33 | 4.08E-21 | 3.03E-20 |
|  | choline | C00114 | 19.633 | 6.97E-10 | 1.26E-09 |
|  | 4-hydroxyphenylacetic acid | C00642 | 104.46 | 7.42E-21 | 5.23E-20 |
|  | 3-indoleacrylic acid | C21283 | 80.881 | 4.85E-19 | 2.50E-18 |

